# The phosphatidylinositol 3-phosphate binding protein SNX4 controls ATG9A recycling and autophagy

**DOI:** 10.1101/2020.06.24.169193

**Authors:** Anthony Ravussin, Sharon A. Tooze, Harald Stenmark

## Abstract

Late endosomes and lysosomes (endolysosomes) receive proteins and cargo from the secretory, endocytic and autophagic pathways. Whereas these pathways and the degradative processes of endolysosomes are well characterized, less is understood about protein traffic from these organelles. In this study, we demonstrate the direct involvement of the phosphatidylinositol 3-phosphate (PI3P) binding SNX4 protein in membrane protein recycling from endolysosomes, and show that SNX4 is required for proper autophagic flux. We show that SNX4 mediates recycling of the transmembrane autophagy machinery protein ATG9A from endolysosomes to early endosomes, from where ATG9A is recycled to the trans-Golgi network in a retromer-dependent manner. Upon siRNA-mediated depletion of SNX4 or the retromer component VPS35, we observed accumulation of ATG9A on endolysosomes and early endosomes, respectively. Moreover, starvation-induced autophagosome biogenesis and autophagic flux were inhibited when SNX4 was downregulated. Altogether, we propose that proper ATG9A recycling by SNX4 sustains autophagy by preventing exhaustion of the available ATG9A pool.

## Introduction

Macroautophagy (hereafter referred to as autophagy) controls numerous fundamental physiological functions [1-3]. During the autophagic process, sequestration of cytoplasmic material ensues within double membrane autophagosomes, which fuse with lysosomes for degradation of contents in the resulting autolysosomes [1, 4, 5]. Conserved autophagy-related gene (ATG) proteins act in a concerted manner in order to regulate proper autophagosome biogenesis and autophagic flux [6]. Strict temporal and spatial regulation of the recruitment of membrane proteins and ATG proteins is required for proper autophagic progression and flux. This is in part achieved by specific membrane lipids and association of lipid-binding proteins, in particular phosphatidylinositol 3-phosphate (PI3P). PI3P is mainly generated by the class III phosphatidylinositol 3-kinase VPS34 through phosphorylation of phosphatidylinositol and has been shown to be essential during the early phase of phagophore biogenesis via recruitment of its effectors DFCP1 and WIPI2 [7, 8]. These proteins are required in order to recruit the only membrane-spanning ATG protein, ATG9A, to the pre-autophagosomal structure and forming phagophore [9, 10].

ATG9A cycles primarily between the Golgi and endosomes in mammalian cells [8] with small amounts found on the plasma membrane [11]. The known residence of ATG9A in endosomal compartments includes EEA1 positive-early endosomes [12], Rab 7-positive late endosomes [13], and Rab11-positive recycling endosomes [14-16]. ATG9A vesicles are essential for autophagosome formation, and studies in yeast have shown that the vast majority of Atg9 vesicles are derived from the Golgi apparatus in a process involving Atg23 and Atg27 and that these vesicles assemble individually into the preautophagosomal structure upon starvation-induced autophagy [17]. In mammalian cells, while the majority of LC3-positive autophagsomes do not contain ATG9A there may be small amounts of ATG9A mislocalized to the membrane of the autophagosome, or indeed arriving from the plasma membrane through late endosomes, which ends up on the limiting membrane of the endolysosome. Although the recruitment and retrieval of ATG9A to and from sites of the forming phagophore is becoming better understood [13, 18], there are still unexplored questions about its trafficking in the endolysosomal system and upon termination of autophagy.

There are multiple PI3P binding proteins other than DFCP1 and WIPI2 expressed in cells, raising the possibility that additional PI3P effectors could be involved in regulation of autophagy. The largest family of PI3P binding proteins is the sorting nexin (SNX) family, a group of membrane-associated proteins containing a phox homology (PX) domain, most of which have been found to bind PI3P [19]. Of the 33 annotated human SNX proteins, a subfamily of SNXs contain a carboxy-terminal BAR domain, and studies of these has shed light on a process of tubular-based endosomal sorting [15]. The SNX-BAR proteins participate in evolutionarily conserved protein complexes that coordinate membrane deformation within the concave surface of dimerization BAR motif which allow association with the phospholipid bilayer through electrostatic interactions possibly for cargo selection [20-22].

In this study, we reveal the involvement of a PI3P binding SNX-BAR protein, SNX4, in ATG9A trafficking and autophagy. We find that SNX4 mediates recycling of ATG9A from endolysosomes and autolysosomes to early endosomes, and that it is essential for proper autophagy.

## Results

### SNX4 localizes to both LAMP1-positive endolysosomes and EEA1-positive early endosomes in a PI3P dependent manner

To determine the localization of SNX4 within the cell, we used SNX4 antibody for detection of endogenous protein and generated a stable retinal pigment epithelial cell line (RPE-1) expressing mNeonGreen-SNX4 for live imaging. Live cell imaging and fixed cell microscopy showed that SNX4 resides on both LAMP1-positive late endosomes/lysosomes (endolysosomes) (23.4% ± 1.2%) and EEA1-positive early endosomes (27.2% ± 3.3%).

SNX4 has been reported to bind PI3P [23] and is implicated in endosomal sorting. We investigated whether PI3P binding is required for recruitment of SNX4 to endosomes and endolysosomes. For this purpose, we incubated cells with SAR405, a highly specific inhibitor of VPS34, which led to rapid loss of SNX4 from endosomes (Movie 1). This shows that SNX4 is recruited to membranes in a PI3P dependent manner.

### ATG9A localizes to Golgi, endolysosomes and early endosomes and partially co-localizes with SNX4

Next, we wanted to determine the relationship of SNX4 to ATG9A. To determine the localization of ATG9A within the cell, we created a double-tagged stable RPE-1 cell line expressing mNeonGreen-SNX4 and mCherry-ATG9A. Live cell imaging showed that many of the SNX4 containing vesicles co-trafficked with ATG9A (Movie 2). Using endogenous antibody staining for ATG9A, we determined that 22.7% ± 6.9 of SNX4 containing vesicles were decorated with ATG9A. Conversely, 41.4% ± 2.3 of ATG9A staining co-localized with SNX4 (Fig. 1G). Additionally, in wild type RPE-1 cells in full medium, we observed a prominent juxtanuclear ATG9A staining which co-localized with the Golgi marker GM-130 (Fig. 1F). 12.5% ± 3.5 ATG9A co-localized with GM130 and 48.9% ± 4.9 Golgi staining contained ATG9A staining. Moreover, 37.0% ± 2.4 of ATG9A co-localized with EEA1-positive early endosomes (Fig. 1D) and 21.4% ± 1.5 of ATG9A co-localized with LAMP1-positive endolysosomes (Fig. 1E). These data confirm that ATG9A is found on both Golgi and endosome membranes, as demonstrated previously [24], and additionally show a significant pool of ATG9A on endolysosomes.

**Figure 1.**
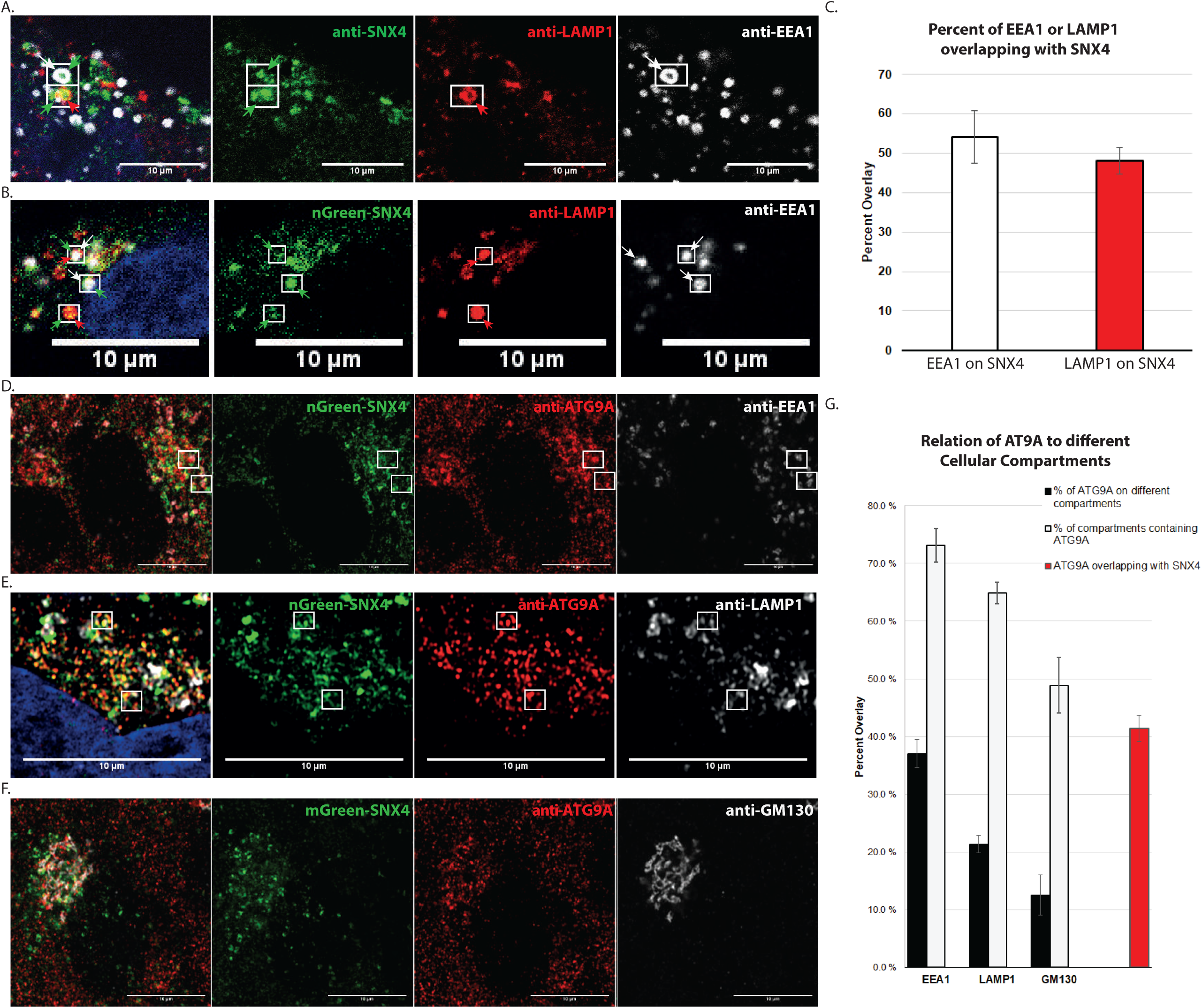
SNX4 co-localizes with both early-endosomal and endolysosomal structures and localizes together with ATG9A on different cellular compartments. A. Endogenous SNX4 co-localizes with both Lamp1 and EEA1 B. mNeonGreen-SNX4n stable cell line co-localizes with both LAMP1 and EEA1 C. Manders Overlay Quantification of overlap between SNX4 and EEA1 and Lamp1 n=3 experiments D. Representative immunofluorescence of ATG9A overlaps with both SNX4 and EEA1 E. Representative immunofluorescence of ATG9A overlaps with both SNX4 and LAMP1 F. Representative immunofluorescence of ATG9A overlaps with both SNX4 and GM130 G. Manders Overlay Quantification of overlap between cellular compartments and ATG9 as well as SNX 4 and ATG9A Means ± SD of n=3-4 experiments with minimum 4 images/experiment

### Inhibition of SNX4 increases the ATG9A localization on endolysosomes upon starvation induced autophagy

As shown in Figure 1, in full medium under basal conditions, ATG9A was found to be mainly juxtanuclear and localizing quite strongly with the Golgi complex. It is thought that this juxtanuclear ATG9A pool traffics through endosomes for fast mobility upon autophagy-inducing stress [13, 14]. Concurrent with these previous studies, we found that in full medium, ATG9A in wild type RPE-1 cells indeed localized to a juxtanuclear region (Fig. 1D,E,F and Fig. 2A). In full medium, SNX4 siRNA did not seem to change the localization of ATG9A, although it slightly increased perinuclear ATG9A intensity. In fact, the portion of LAMP1 vesicles containing ATG9A in control compared to SNX4 knockdown cells was no different (64.3% ± 3.3 and 65.5% ± 2.0, control and SNX4siRNA respectively) (Fig. 2A and D).

**Figure 2.**
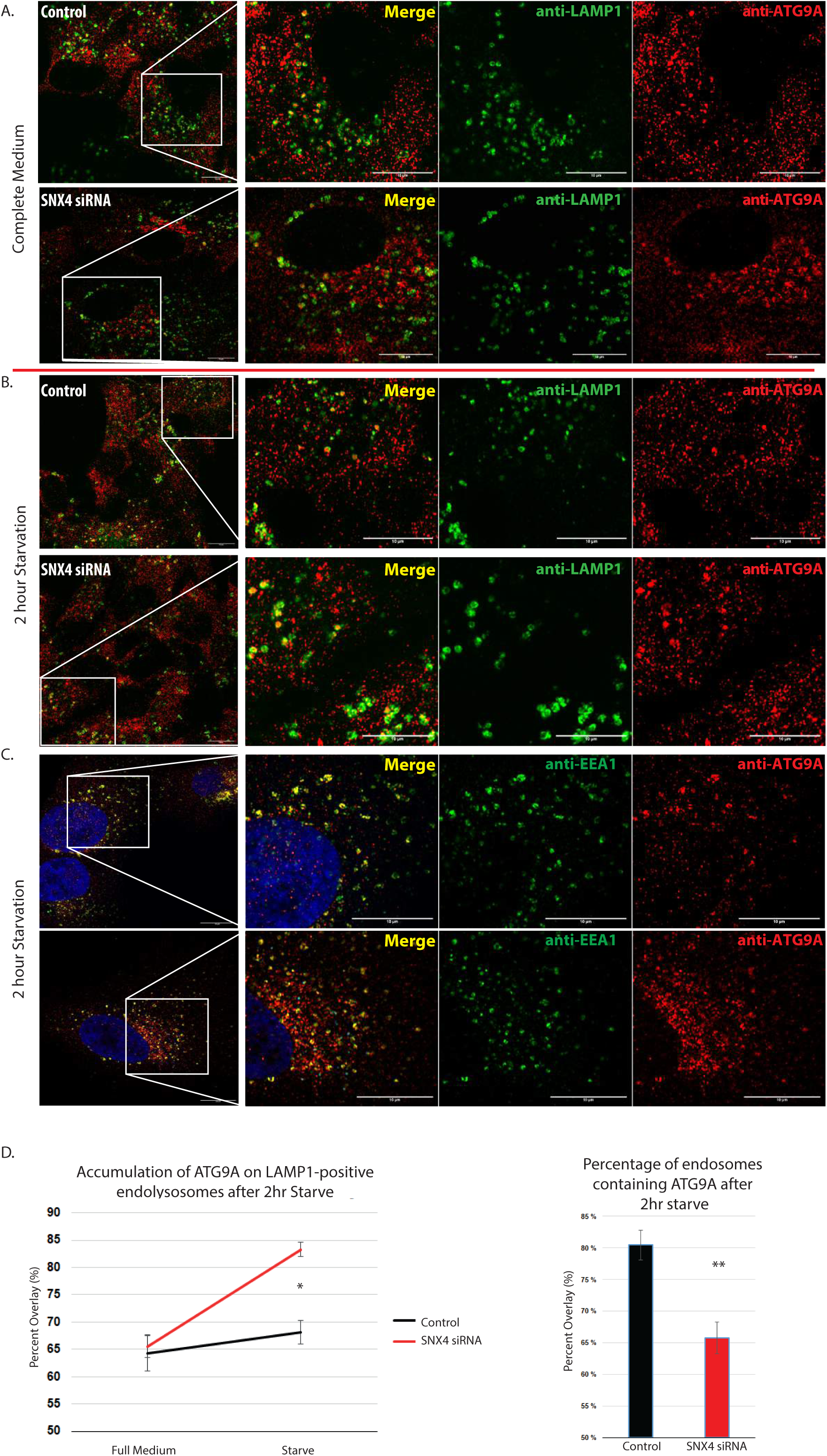
ATG9A increases on LAMP1 positive structures and decreases on EEA1 positive structures during autophagy in SNX4 depleted cells. A. Representative immunofluorescence image of ATG9A overlapping with LAMP1 in full medium B. Representative immunofluorescence image of ATG9A overlapping with LAMP1 after 2 hours starvation C. Representative immunofluorescence image of ATG9A overlapping with EEA1 after 2 hours starvation D. In starved conditions, ATG9A increases co-localization with LAMP1 positive structures upon SNX4 depletion. Manders overlay quantifications Means ± SD of N>3 experiments

Next, we wanted to assess whether ATG9A localization changes upon inhibition of SNX4 when autophagy was induced by amino acid starvation. Strikingly, upon SNX4 knockdown, ATG9A was redistributed to a more peripheral localization (Fig. 2B). Consistent with previous studies, starvation induced autophagy in wild type RPE-1 cells also triggered dispersal of the ATG9A Golgi pool into a more peripheral dispersion. Interestingly, upon starvation ATG9A did not seem to disperse and had a similar juxtanuclear localization in SNX4 siRNA treated cells when compared to controls (Fig. 2B). Upon quantification, ATG9A was found to accumulate in LAMP1-positive endolysosomes in SNX4 siRNA inhibited cells upon starvation (Fig. 2D) potentially leading to larger LAMP1 vesicles (Fig. 2B). The percentage of LAMP1 positive vesicles containing ATG9A increased in SNX4 siRNA cells to 83.2% ± 1.3 (Fig. 2D). Furthermore, ATG9A co-localization decreased with the early-endosomal marker EEA1 (Fig. 2E). These data suggest that SNX4 is required for precise mobilization of the ATG9A protein during autophagy.

### Inhibition of retromer increases the localization of ATG9A on early endosomes

VPS35 is the core functional component of retromer complex variants active to ensure proper sorting of selected transmembrane cargo proteins from endosomes to the biosynthetic pathway. VPS35 is recruited to endosomal membranes and with other SNX-BAR retromer, mediates retrograde transport of cargo proteins from endosomes to the trans-Golgi network (TGN) [25]. Previous studies have shown no effect of VPS26 retromer in ATG9A redistribution or in the reestablishment of ATG9A juxtanuclear population [18]. To investigate whether ATG9A is recycled back to the Golgi via retromer complex in RPE-1 cells, we used siRNA inhibition of VPS35 to determine the fate of ATG9A. Interestingly, upon VPS35 knockdown, we observed that ATG9A had a stronger juxtanuclear localization (Fig. 3A and B). However, the peripheral ATG9A in these VPS35 depleted cells showed an increased co-localization with the early-endosomal marker EEA1 (Fig. 3D). The localization of ATG9A to EEA1 positive endosomes increased from 71.9% ± 0.7 in control to 80.9% ± 1 in VPS35 depleted cells (Fig. 3D). This confirms the importance of retromer in recycling of ATG9A recycling from endosomes to the Golgi.

**Figure 3.**
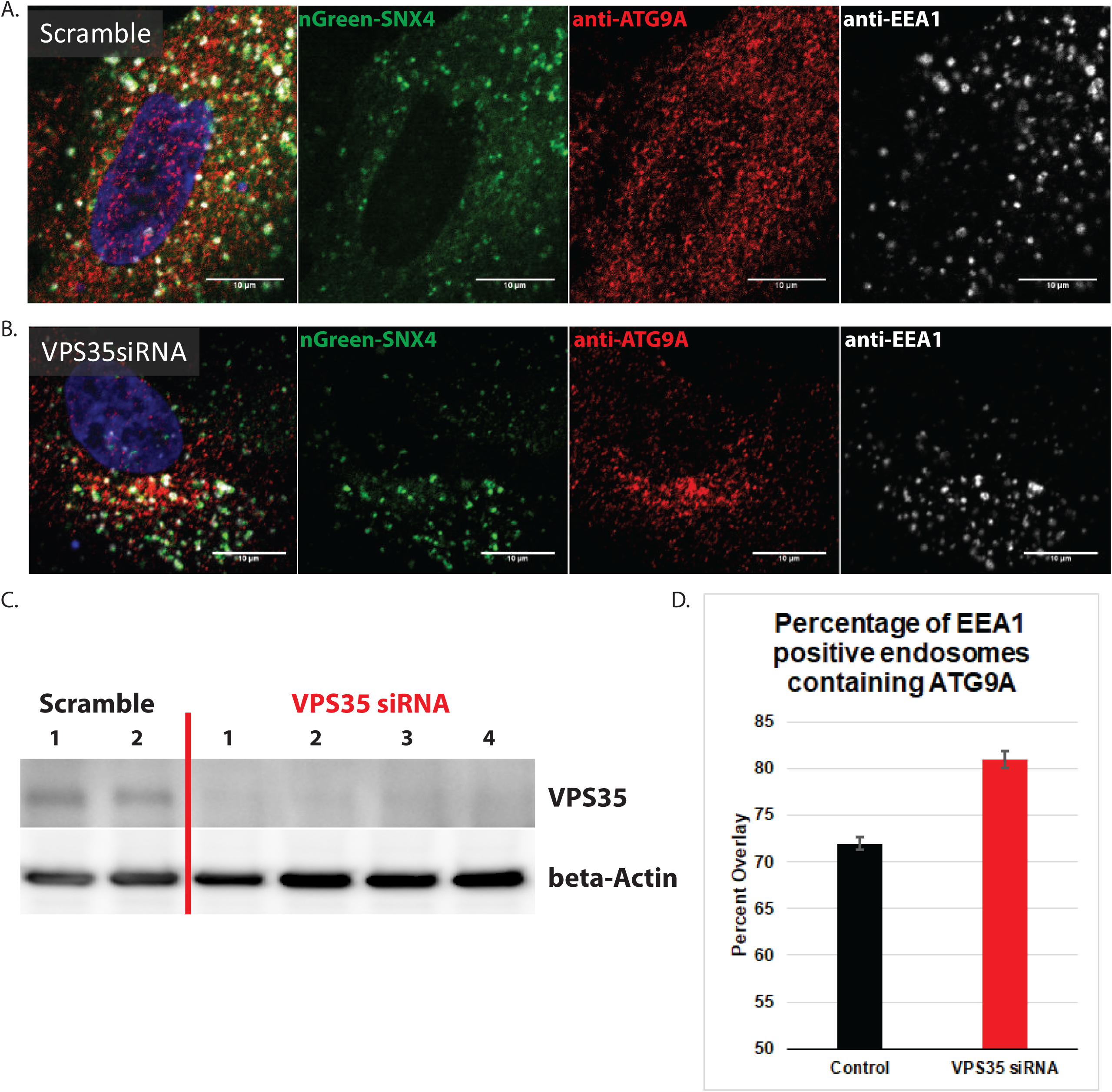
ATG9A increases on early endosomes during autophagy when VPS35 is depleted. A. Representative immunofluorescence of ATG9A overlapping with both SNX4 and EEA1 in scramble control RPE-1 cells B. Representative immunofluorescence of ATG9A overlapping with both SNX4 and EEA1 in VPS35 depleted RPE-1 cells C. Western blot expressing VPS35 knockdown efficiency D. Manders Overlay Quantification of overlap between ATG9A and EEA1 n=2 experiments (5 technical replicates per experiment)

### SNX4 is required for formation of LC3-positive autophagosomes upon starvation

To examine whether SNX4 is necessary for proper autophagy in mammalian cells, we tested the necessity of SNX4 during amino acid starvation induced autophagy. During autophagy, the cytosolic form of Microtubule-associated protein 1A/1B-light chain 3 (LC3), LC3-I, is conjugated to phosphatidylethanolamine (PE) to form LC3-PE conjugate (LC3-II), which is recruited to autophagosomal membranes. Using an imaging based approach, we observed that starvation-induced LC3 structures were significantly decreased upon inhibition of SNX4 (Fig. 4A). Quantitative image analyses showed that there were decreases in average LC3 area, average size of LC3 as well as total LC3 intensity per cell (Fig. 4C).

**Figure 4.**
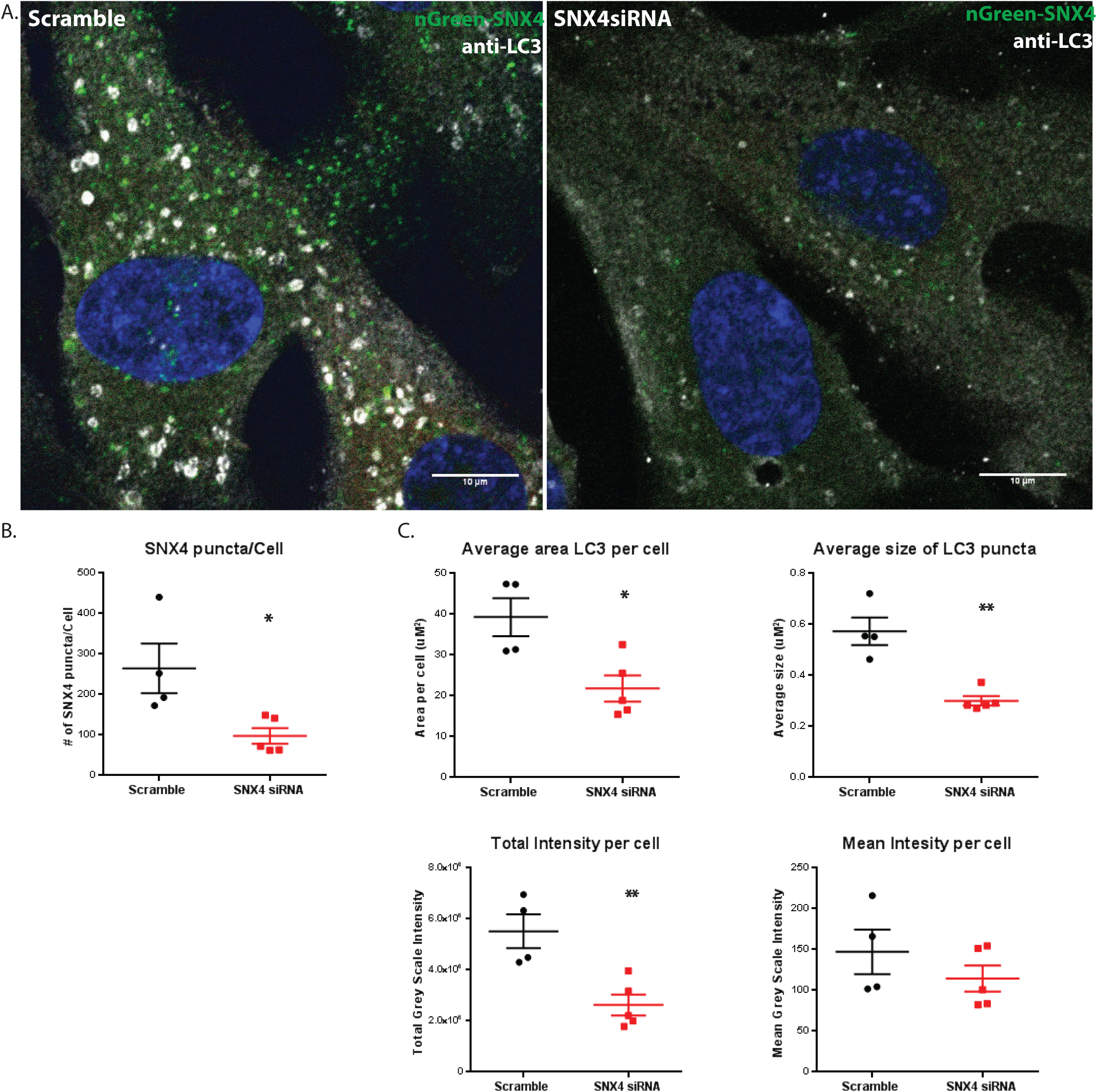
Depletion of SNX4 inhibits starvation-induced formation of LC3-positive autophagosomes. A. Representative immunofluorescence micrograph of LC3 upon starvation in scramble (left) and SNX4 depleted (right) RPE-1 cells. B. Quantifications of SNX4 puncta in scramble and SNX4 depleted RPE-1 cells C. Quantifications of area of LC3 and average area of LC3/cell, size of LC3 puncta, total intensity/cell, and mean intensity/cell in scramble and SNX4 siRNA All quantifications n=3 experiments with 2 technical replicates in scramble and 3 replicates in SNX4 siRNA cells.

### Depletion of SNX4 increases the ATG9A localization on autolysosomes upon starvation induced autophagy

To assess if the ATG9A increase on late endosomes and lysosomes are related to a decrease in autophagy, we investigated the presence of ATG9A on autolysosomes, characterized by their content of both LC3 and LAMP1. In basal, full medium conditions, we again observed the juxtanuclear ATG9A staining. In RPE-1 control cells, ATG9A was present on 54.5% ± 1.8 of LC3-positive, LAMP1-positive autolysosome structures (Fig. 5A and C). Upon 2 hour starvation in control cells, there was no increase of ATG9 on autolysosomes, 55.2% ± 2.4 (Fig. 5B and C). In SNX4 siRNA depleted cells, there was a marked decrease in the total number of LC3 positive structures, consistent with the findings in Fig. 5. Nevertheless, ATG9A increased the relative localization with autolysosomes (LC3+,LAMP1+) upon amino acid starvation (Full medium, 53.6% ± 2.0 and Starvation, 67.1% ± 2.2) (Fig. 5 A,B,C). This relative increase of ATG9A on autolysosomes in cells lacking SNX4 is presumably due to the inability of ATG9A to recycle from the autolysosomes for reutilization in another round of autophagy.

**Figure 5.**
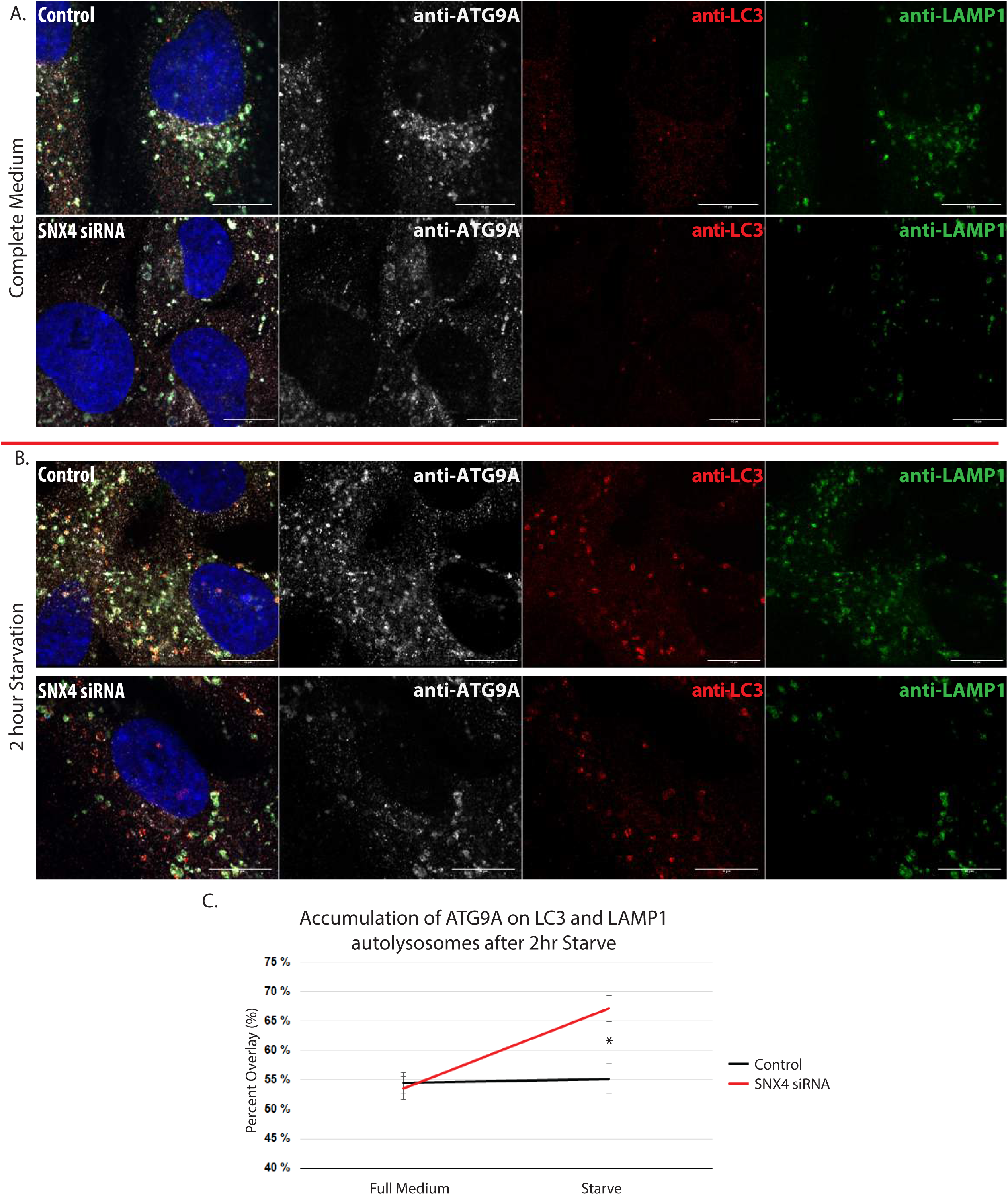
In SNX4 depleted cells, ATG9A increases on LC3- and LAMP1-positive autolysosomes. A. Representative immunofluorescence micrograph of ATG9A overlapping with both LC3 and LAMP1 in full medium B. Representative immunofluorescence micrograph of ATG9A overlapping with both LC3 and LAMP1 after 2 hours starvation C. Manders overlay quantifications. Under starved conditions, ATG9A increases co-localization with autolysosomes upon SNX4 depletion. n= 3 experiments

### SNX4 is required for proper autophagic flux

To quantitatively assess the effect of SNX4 on bulk autophagic flux, we performed a long-lived protein degradation assay [26]. This pulse-chase labelling approach showed that inhibiting SNX4 in amino acid starved cells decreased the total protein degradation by 2.4% ± 0.2, 2.7% ± 0.2 and 2.5% ± 0.2 (three separate experiments) when compared to wild type. Effectively, this showed that cells lacking SNX4 had a 44.4% protein degradation when compared to control cells in full medium (Fig. 6A) and a 44.9% protein degradation when compared to the wild type upon starvation (Fig. 6B). Taken together, these data show that SNX4 is required for proper autophagic flux.

**Figure 6.**
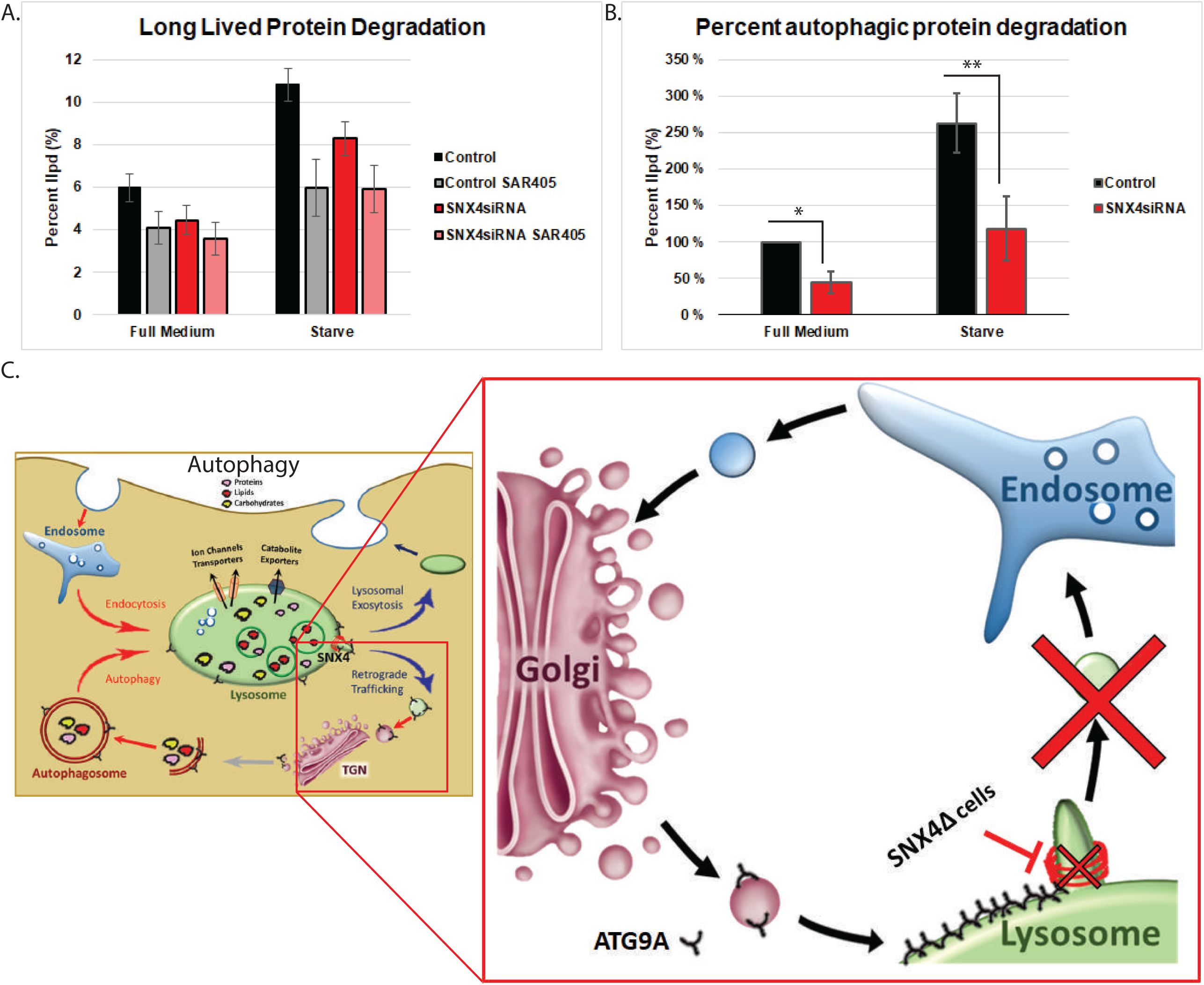
SNX4 depletion prevents proper autophagic flux. A. Quantification of the percent total degraded long-lived proteins (Dark bands) and proteosomal long lived degraded proteins, cells treated with SAR405 (opaque bands). n=3 different experiments B. Quantification of the percent change in autophagic long lived protein degradation (p=0.049 and p=0.002) n=3 experiments normalized to percentage of control proteins degraded C. Graphical Model of SNX4 dependent ATG9A recycling from autolysosomes to endosomes

## Discussion

Whereas PI3P has a well-described function in protein recruitment to the growing phagophore, we have here uncovered a new role of this lipid in autophagy. We have used microscopy of live and fixed cells to show the requirement of the PI3P effector SNX4 for proper autophagic flux and for recycling of the transmembrane autophagy protein ATG9A from endolysosomes and autolysosomes. It has been observed that Snx4 can bind phosphatidylserine in budding yeast [27], but the punctate localization of Snx4 is dependent on the presence of PI3P [28]. Mammalian SNX4 is PI3P dependent as observed with its disassociation from vesicles upon Class III phosphatidylinositol 3-kinase VPS34 inhibition by SAR405. We propose that recycling of ATG9A from endolysosomes and autolysosomes sustains autophagy by allowing the same ATG9A molecule to be used for multiple cycles of vesicle delivery (Fig. 6C).

ATG9A vesicles are mobile and their trafficking is controlled via nutrient regulated signals [13]. Previous studies have found the importance of ATG9A during autophagosome formation, proposedly by functioning in vesicular delivery to the phagophore initiation site [18]. Upon starvation-induced autophagy, ATG9A redistributes from the perinuclear Golgi regions to an enrichment juxtaposed to the autophagosome initiation site [13, 14, 16, 29]. Because ATG9A has been shown to be involved in the early steps of autophagy, we suggest that recycling of ATG9A from the endolysosome back to the Golgi is required to sustain proper autophagy.

SNX proteins have been previously implicated in tubular sorting of endocytic proteins and involved in endosome to Golgi retrograde transport of ricin [30]. Interestingly, binding of another PX-BAR domain containing protein, SNX18, to PI(4,5)P_2_ is required for regulation of autophagy via control of ATG9A trafficking from recycling endosomes and formation of ATG16L1- and WIPI2-positive autophagosome precursor membranes [15]. Thus, cycling of membrane proteins via SNX-BAR proteins is necessary for proper autophagy. In the present study, we show a co-localization of SNX4 and ATG9A in several cellular compartments. When SNX4 is inhibited, ATG9A is unable to traffic correctly and remains in a more juxtanuclear Golgi zone. Although this is true in full medium conditions, this is even more pronounced during starvation-induced autophagy.

We found that, in RPE-1 cells lacking SNX4, proper autophagic responses are altered as revealed by several independent autophagy assays, including fluorescence microscopy and long-lived protein degradation assays. During starvation-induced autophagy, there was a decrease in LC3 puncta as well as long-lived protein degradation. Even though our data point to a role of SNX4 in ATG9A recycling, it is also conceivable that SNX4 regulates autophagy through additional mechanisms. Evidence from budding yeast suggests that Snx4 promotes autophagy and vacuole membrane fusion. In cells lacking Snx4 and starvation induced, phosphatidylserine was shown to accumulate in the membranes of the endosome and vacuole, autophagy intermediates accumulated within the cytoplasm, and homotypic vacuole fusion was impaired. The Snx4-Atg20 dimer displayed preference for binding and remodeling of phosphatidylserine-containing membrane in vitro, which suggested that Snx4-Atg20-coated carriers export phosphatidylserine-rich membrane from the endosome [27]. Whether SNX4 is directly required for maintaining glycerophospholipid homeostasis and autophagy/vacuole fusion, remains to be investigated in mammalian cells.

Overall, the data here within provide a novel recycling process of integral membrane proteins from endolysosomes in mammalian cells, similar to protein recycling from the vacuole in budding yeast [31]. During autophagy, SNX4 is recruited to endolysosomes via its PI3P binding PX domain and is required for recycling of ATG9A. In the absence of SNX4, ATG9A and presumably additional unidentified proteins, are unable to be recycled back to the Golgi for utilization in additional rounds of autophagy. Under stress, this inability to properly recycle ATG9A prevents proper autophagic flux from occurring. Overall, our findings indicate that PI3P is not only important in the initial stages of autophagy but also for the late stage of autophagy through recycling of ATG9A.

## Materials & Methods

### Cell Culture and generation of stable cell lines

hTERT-RPE-1 cells (human retinal pigment epithelial cells immortalized with telomerase) and stable cell lines derived from these cells were maintained in DMEM-F12/Dulbecco’s Modified Eagle’s Medium high glucose (DMEM, Sigma-Aldrich, D0819) supplemented with 10% fetal bovine serum (Sigma Aldrich, F7524), 100 U/ml penicillin and 100 μg/ml streptomycin. All cells were cultured in a humidified incubator at 37°C supplemented with 5% CO2. For amino acid and growth factor starvation experiments, the growth medium was removed, cells washed 3 times with EBSS (GIBCO BRL, 24,010–043) and replaced with EBSS or Live Cell Imaging Solution (Molecular Probes, A14291DJ), supplemented with 20 mM glucose (Merck, 108,342).

Both stable cell lines used in these studies were lentivirus-generated pools, using plasmids (described below) pcDH-PGK-mNeonGreen-SNX4-IRES-Puro and pCDH-PGK-mCherry-ATG9A-IRES-Neo. The weak PGK promoter was used for transgene expression at rather low expression levels. Third generation lentivirus was generated as previously published in [32]. Briefly, mCherry and mNeonGreen fusions were generated as Gateway ENTRY plasmids using standard molecular biology techniques. From these vectors, lentiviral transfer vectors were generated by recombination into customized pCDH (System Biosciences CD532-A) destination vectors using a Gateway LR reaction. VSV-G pseudotyped lentiviral particles were packaged using a third-generation packaging system that was a gift from Didier Tromo (deposited by Tromo at Addgene, 12,251, 12,253 and 12,259). Cells were then transduced with low virus titers and stable expressing populations were generated by antibiotic selection.

### Antibodies

The following antibodies were used for the studies: Hamster anti-ATG9A ([33], Immunofluorescence 1:500), Rabbit anti-LAMP1 from Sigma-Aldrich (L1418, Immunofluorescence 1:200), Mouse anti-LAMP1 from BD (555798, Immunofluorescence 1:500), Human anti-EEA1 provided by Ban-Hock Toh (Monash University, Immunofluorescence 1:160,000), Mouse anti-GM130 from BD (610823, Immunofluorescence 1:250), Sheep anti-TGN46 from Biorad (AHP500, Immunofluorescence 1:500), Hoechst 33342 from Invitrogen Molecular Probes (H3570).

### Immunostaining

Cells grown on coverslips were fixed with 4% formaldehyde (Polyscience, 18814) for 20⍰min in room temperature and permeabilized with 0.05% Saponin (Sigma-Aldrich, S7900) or 0.05% Digitonin (Sigma-aldrich, D141) in PBS for 5 minutes. Cells were then blocked in 5% BSA for 20 minutes and stained with the indicated primary antibody concentrations for 1⍰hour in 1%BSA. The coverslips were then washed in PBS/Saponin for 5 minutes and stained for 1⍰hour with fluorescently labeled secondary antibody at 1:1000 concentration in dark. The cells were then washed with PBS and water and were mounted using mowiol (Sigma-Aldrich) alone or supplemented with Hoechst.

### siRNA transfections

Silencer Select siRNAs against human SNX4 and VPS35, and nontargeting control “scrambled” siRNA (predesigned, 4,390,844) were purchased from Ambion^®^ (Thermo Fisher Scientific). Cells at 30% confluency were transfected with 20 nM final siRNA concentration using Lipofectamine RNAiMax transfection reagent (Life Technologies, 13,778–150) according to the manufacturer’s instructions and used for experiments after 48 hours for VPS35 or 72 hours for SNX4. All siRNA oligonucleotides have been validated previously for target specificity and knockdown levels were routinely confirmed by western blotting or immunofluorescence imaging.

### Immunoblotting

Cells were washed in cold PBS and lysed in 2x Laemmli Sample Buffer (Bio-Rad Laboratories, 1,610,737) supplemented with DDT. Whole-cell lysates were separated on SDS-PAGE on 4–20% gradient gels (mini-PROTEAN TGX; Bio-Rad). Proteins were transferred to polyvinylidene difluoride (PVDF) membranes (TransBlot^®^ TurboTM LF PVDF, Bio-Rad) followed by 1 hour blocking in 3% BSA and overnight antibody incubation in Tris-buffered saline with 0.1% Tween-20 (Sigma-Aldrich, P1379) at 4 degrees. Membranes incubated with HRP (horseradish peroxidase)–conjugated antibodies (HRP rabbit, 111 035 144; HRP mouse, 115 035 146) were developed using Clarity western ECL substrate solutions (Bio-Rad) with a ChemiDoc XRS+ imaging system (Bio-Rad).

### Confocal Fluorescence Microscopy

Confocal images were obtained using LSM 880 with Airyscan confocal microscope (Carl Zeiss) equipped with laser lines: 405, 458, 488, 514, 561, and 633 nm and objectives 10x NA 0.45 DIC II (Plan-Apochromat), 20x NA 0.8 DIC II (Plan-Apochromat), 25x NA 0.8 (LD LCI Plan-Apochromat), 40x NA 1.2 Water Imm DIC III (C-Apochromat), 63x NA 1.4 oil DIC III (Plan-Apochromat). Images were acquired using the 63x/1.40 oil DIC III.

### Image processing and data analysis

hTERT-RPE-1 cells stably expressing mNeonGreen-SNX4 were fixed and stained with different antibodies. Images were acquired by concocal fluorescence microscopy fixed intensity below saturation. Colocalization was then quantified with ImageJ/FIJI (https://imagej.net/Fiji) using the JACoP plugin [34]. Manders’ Overlap Coefficient was used to describe the amount overlap [35].

### Statistical analysis

Statistical analysis was performed using Graphpad Prism. Student’s two-tailed t test was used to measure for statistical significance in samples with a gaussian distribution. In order to account for differences in staining efficiencies and imaging conditions, experiments involving quantification of intensities were normalized by the mean of the experiment and then analyzed. This is also true of the percent long-lived protein degradation experiments. *P⍰< ⍰0.05, **P⍰< ⍰0.01.

## Acknowledgments

We thank Kia-Wee Tan for help with the construct design of plasmids and SNX4 cell lines. The Core Facilities for Advanced Light Microscopy and Advanced Electron Microscopy at Oslo University Hospital are acknowledged for access to relevant microscopes. Minoo Razi is acknowledged for help with imaging experiments. We thank all the members of the Molecular Cell Biology Stenmark Laboratory for comments and helpful discussion. This work was supported by the Norwegian Cancer Society (project number 182698), the South-Eastern Norway Regional Health Authority (grant number 2018081), the European Research Council (grant number 788954), and the Research Council of Norway (project number 262652) through its Centres of Excellence funding scheme. This work was also partly supported by a STSM Grant from COST Action CA15138 (TRANSAUTOPHAGY). S.A.T was supported by The Francis Crick Institute which receives its core funding from Cancer Research UK (*FC001187*), the UK Medical Research Council (*FC001187*), and the Wellcome Trust (*FC001187*).

